# Double-strand breaks can induce DNA replication and damage amplification in G2 phase-like oocytes of mice

**DOI:** 10.1101/2020.06.29.179010

**Authors:** Jun-Yu Ma, Xie Feng, Feng-Yun Xie, Sen Li, Lei-Ning Chen, Shi-Ming Luo, Xiang-Hong Ou

## Abstract

Break-induced DNA replication (BIR) have been detected not only in the genome of rare disease patients but also in cancer cells, however, the mechanisms of BIR formation haven’t been explained in details. In the late G2 phase-like mouse oocytes, we found DNA double-strand breaks (DSBs) could induce Rad51 dependent small-scale DNA replication. In addition, we also found the DSBs could be amplified in mouse oocytes, and the amplification could be inhibited by Rad51 inhibitor IBR2 and DNA replication inhibitor ddATP. Lastly, we found the DSB repair was relatively inefficiency in hybrid mouse oocytes compared with that of the purebred mouse oocytes. We found DSBs could induce BIR more easier in hybrid mouse oocytes, indicating the DNA repair in oocytes could be affected by the sequence differences between homologous chromatids. In summary, our results indicated that the condensed chromatin configuration in late G2 phase and the sequence similarity between broken DNA and template DNA are causing factors of BIR in mammalian genome, and the DNA damage could be amplified in late G2 phase cells.

## Introduction

DNA double-strand breaks (DSBs) in cells could be repaired by multiple pathways like homologous recombination (HR), non-homologous end joining and single-strand annealing (Ceccaldi *et al.* 2016). Generally, during the HR repair of DSBs, if the broken DNA end invaded into the allelic sequence, the repair process would be mediated by the mechanism of synthesis-dependent strand annealing (SDSA)(Miura *et al.* 2012) or double-holiday junctions (dHJ)(Bzymek *et al.* 2010). If the broken DNA end invade into a non-allelic sequence, it might initiate the non-allelic homologous recombination (NAHR) pathway which would cause recurrent copy number variant (CNV) formation (Inoue and Lupski 2002; Gu *et al.* 2008; Liu *et al.* 2012). If the broken DNA end had invaded into a non-allelic sequence but the synthesized new DNA single strand couldn’t form dHJ structure or reanneal with the other broken end, then it might induce break-induced replication (BIR) (Malkova and Ira 2013) or even template switching (Lee *et al.* 2007; Hastings *et al.* 2009; Anand *et al.* 2014; Li *et al.* 2020) in genome, and both BIR and template switching could induce complex genomic rearrangement (CGR) in the genome (Zhang *et al.* 2013; Pellestor and Gatinois 2018).

CGR is a common somatic structure variation pattern in cancer cells (Sudmant *et al.* 2015; Li *et al.* 2020). CGR has also been found formed *de novo* in germline cells (Kloosterman *et al.* 2011), which might induce rare diseases. Based on the genomic data of human family trios, the specific germline CGRs had been proposed to be formed at the peri-zygotic period and depend on the microhomology-mediated BIR (mmBIR) pathway (Liu *et al.* 2017). However, the mechanisms of how CGR is formed in peri-zygotic cells, such as oocytes, spermatocytes and/or early embryos, haven’t been widely studied yet.

For mammalian females, their oocytes finish meiotic HR at fetal stage and arrest at the G2 phase-like dictyate stage before or after their born (MacLennan *et al.* 2015; Dalbies-Tran *et al.* 2020). Then the arrest of G2 phase-like stage oocytes will be maintained for weeks or even tens of years according to different species. When the female mammals are sex-matured, their oocytes can be activated and start to grow. For mouse, oocyte growth will take about two weeks to accumulate maternal materials for subsequent embryo development (Gosden *et al.* 1997; Li *et al.* 2010). When mouse oocytes are fully grown, their transcription activities will be silenced and their DNA will be condensed and form a ring like Hoechst positive structure Surround the Nucleolus (Dumdie *et al.* 2018). These fully grown oocytes are termed SN oocytes whereas the growing oocytes are termed non-SN (NSN) oocytes (Tan *et al.* 2009). For the NSN and SN oocytes, both endogenous metabolites and exogenous factors can induce DSBs in their nuclear genome (Carroll and Marangos 2013; Tubbs and Nussenzweig 2017; Winship *et al.* 2018; Stringer *et al.* 2020), but whether DSBs in growing NSN and fully grown SN oocytes could be the contributing factor of CGR hasn’t be analyzed. Our previous works showed that exogenous DSBs induced the chromatin in SN oocytes to be entangled and matted together by Rad51 (Ma *et al.* 2019a), in this study we further analyzed the features of DSB repair in oocytes which would be new clues of CGR formation in germ cells and somatic cells.

## Results

### DSBs induce DNA replication in SN oocytes

To analyze the DSB repair in growing and fully grown oocytes, NSN and SN stage oocytes from large antral follicles were treated with 10 μM Bleomycin for 1 hour. To determine whether DSBs could induce DNA replication, 10 μM EdU was added into the culture media. After Bleomycin treatment, oocytes were recovered in Bleomycin-free media for 15 hours. After that oocytes were fixed and EdU were marked by the click reaction (Hein *et al.* 2008). As a result we could find visible EdU signals in the nuclear of NSN-SN and SN oocytes but not in nuclear of NSN oocytes and control oocytes (Figure 1A), indicating DSB could induce BIR in NSN-SN and SN oocytes.

**Figure 1.**
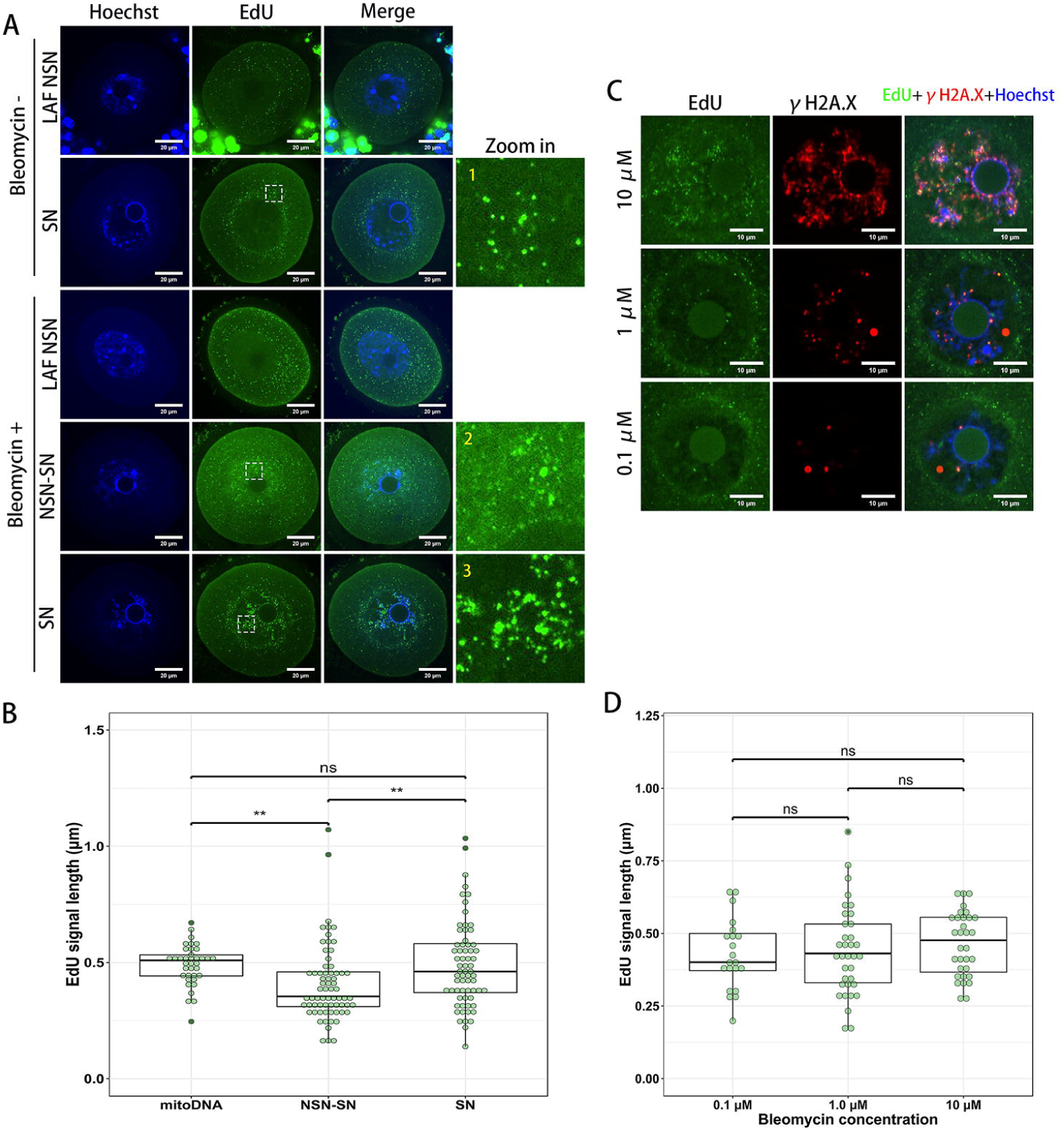
DNA DSBs induced small-scale BIR in NSN-SN and SN stage oocytes but not NSN oocytes. (**A**) Oocytes are treated with or without 10 μM Bleomycin for 1 hour and released from Bleomycin for 15 hours. EdU signals (green) could be seen in Belomycin treated NSN-SN oocytes and SN oocytes. (**B**) The EdU signal sizes in SN stage oocytes are larger than NSN-SN oocytes but comparable to mtDNA replication induced EdU signals. (**C** and **D**) EdU sizes induced by different Bleomycin doses have no significant difference. (**C**) γH2A.X foci can be observed beside to or overlapped with the EdU signals. LAF, large antral follicle.

As there were persistent mitochondrial DNA (mtDNA) replication events in NSN and SN oocytes (Figure S1), so we compared the EdU signal sizes in the nuclear of NSN-SN oocytes and SN oocytes with the mtDNA replication induced EdU sizes. We found the EdU signal sizes in SN oocyte nuclear and mitochondria were comparable, however, the EdU signal sizes in NSN-SN oocyte nuclear were less than that in SN oocytes and mitochondria (Figure 1B).

By immunofluorescence labeling of the DSB marker γH2A.X, we found most nuclear EdU signals were connected with or adjacent to the γH2A.X foci (Figure 1C). When we induce the DSB with different Bleomycin dose, the γH2A.X focus and EdU signal number decreased obviously as the Bleomycin concentration decrease, but the EdU sizes had no significant difference (Figure 1D).

### BIR in oocytes is Rad51 dependent

After 1 μM Bleomycin treatment for 1 hour, we recovered the oocytes in Bleomycin-free media for 0 hour, 12 hours or 24 hours. By counting the γH2A.X foci number in different group oocytes we found γH2A.X foci numbers decreased in recovered oocytes as time extension (Figure 2A), indicating the DSBs could be repaired in SN oocytes. Similar with our previous results (Ma *et al.* 2019a), we found HR protein Rad51 colocalized with γH2A.X in oocytes which were treated with 1 μM Bleomycin and recovered for 2 hours, 12 hours or 24 hours (Figure 2B). So it’s not surprising that when we marked out the EdU signals in DSB SN oocytes, the EdU signals were adjacent to the Rad51 foci (Figure 2C). Although we could also detect Rad51 foci in DSB NSN oocytes, however, there was no detectable EdU signal in these NSN oocytes (Figure 2C).

**Figure 2.**
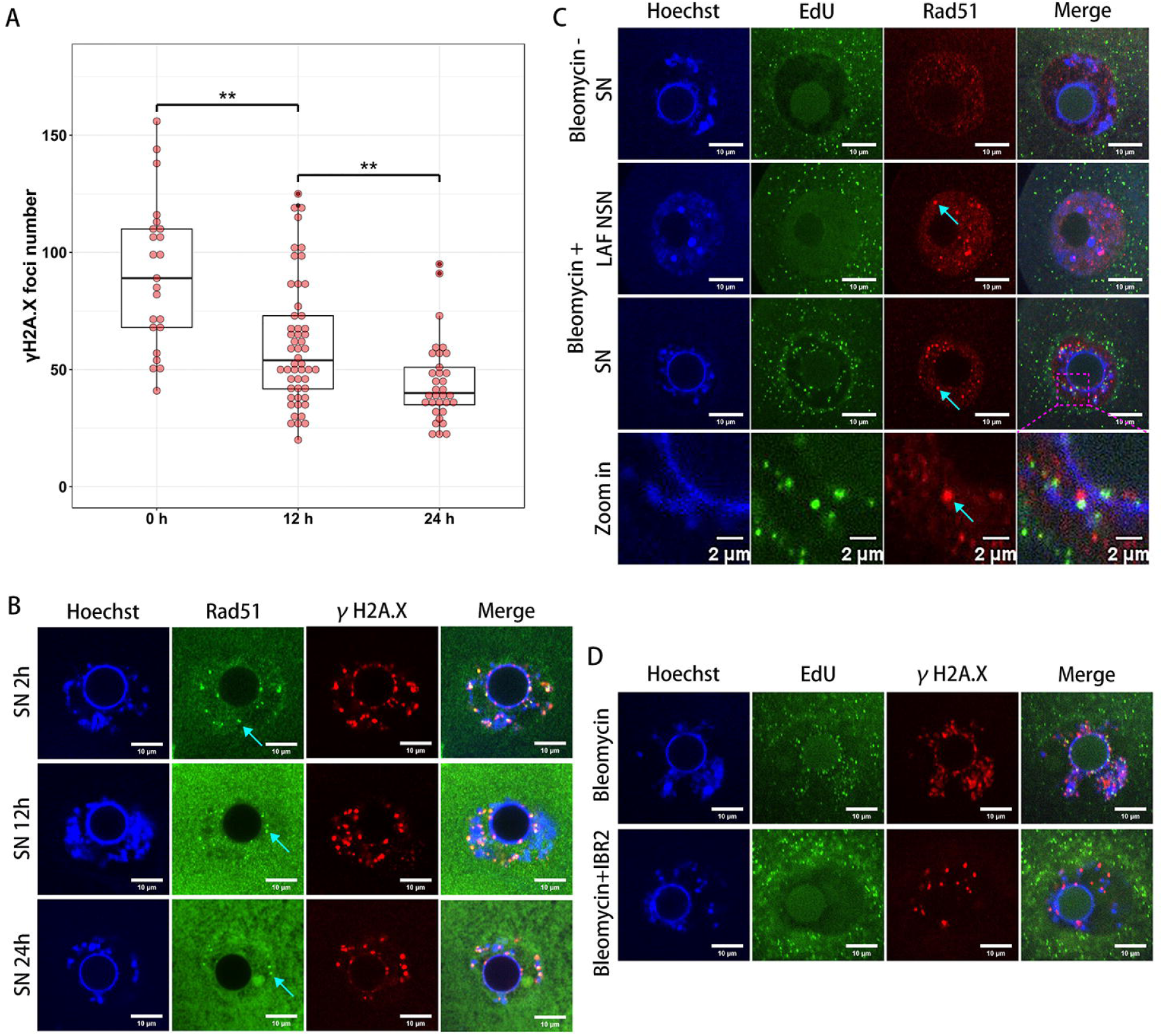
Rad51 plays essential role in BIR initiation in DSB oocytes. (**A**) DSBs could be repaired in SN stage oocytes. γH2A.X foci in oocytes are counted 0, 12, or 24 hours after 1 hour treatment of 1 μM Bleomycin. (**B**) Rad51 foci can be detected overlapped with γH2A.X foci. (**C**) Rad51 foci are relate to the EdU signals. (**D**) When treated oocytes with Rad51 inhibitor IBR2 (100 μM) for 5 hours and then introduced DSBs with 1 μM Bleomycin for 1 hour and released oocytes for 12 hours, we could find the formation of EdU signals are suppressed. Rad51 foci are marked by arrows. LAF, large antral follicle.

To analyze whether the DSB induced oocyte BIR was mediated by Rad51, we treated the oocytes with or without the Rad51 inhibitor IBR2 (Zhu *et al.* 2015). We first treated the oocytes with or without 100 μM IBR2 for 5 hours and then these oocytes were treated with 1 μM Bleomycin for 1 hour. Then the oocytes were recovered from Bleomycin for 15 hours. After that oocytes were fixed and we marked EdU and γH2A.X by click reaction and immunofluorescence labeling. As a result we found the BIRs were obviously suppressed in IBR2 treated oocytes (Figure 2D), indicating the BIR in DSB SN oocytes is Rad51 dependent.

### DSBs could be amplified in SN oocytes

It had been reported that RAD51 could mediate a multi-invasion process in yeast which can amplify the initial DNA DSB numbers in cells (Piazza *et al.* 2017). To test whether Rad51 filaments in DSB SN oocytes could amplify the initial DSBs, we measured the γH2A.X signal levels in DSB oocyte treated with or without IBR2. We first treated oocytes with 100 μM IBR2 for 5 hours and induce DSBs using 1 μM Bleomycin. When DSB oocytes were recovered from Bleomycin for 4 hours, we found the γH2A.X signals were decreased significantly in IBR2 treated oocytes comparing to that in control oocytes (Figure 3A and 3B), indicating the DSBs in oocytes were indeed amplified and the amplification process was mediated by Rad51.

**Figure 3.**
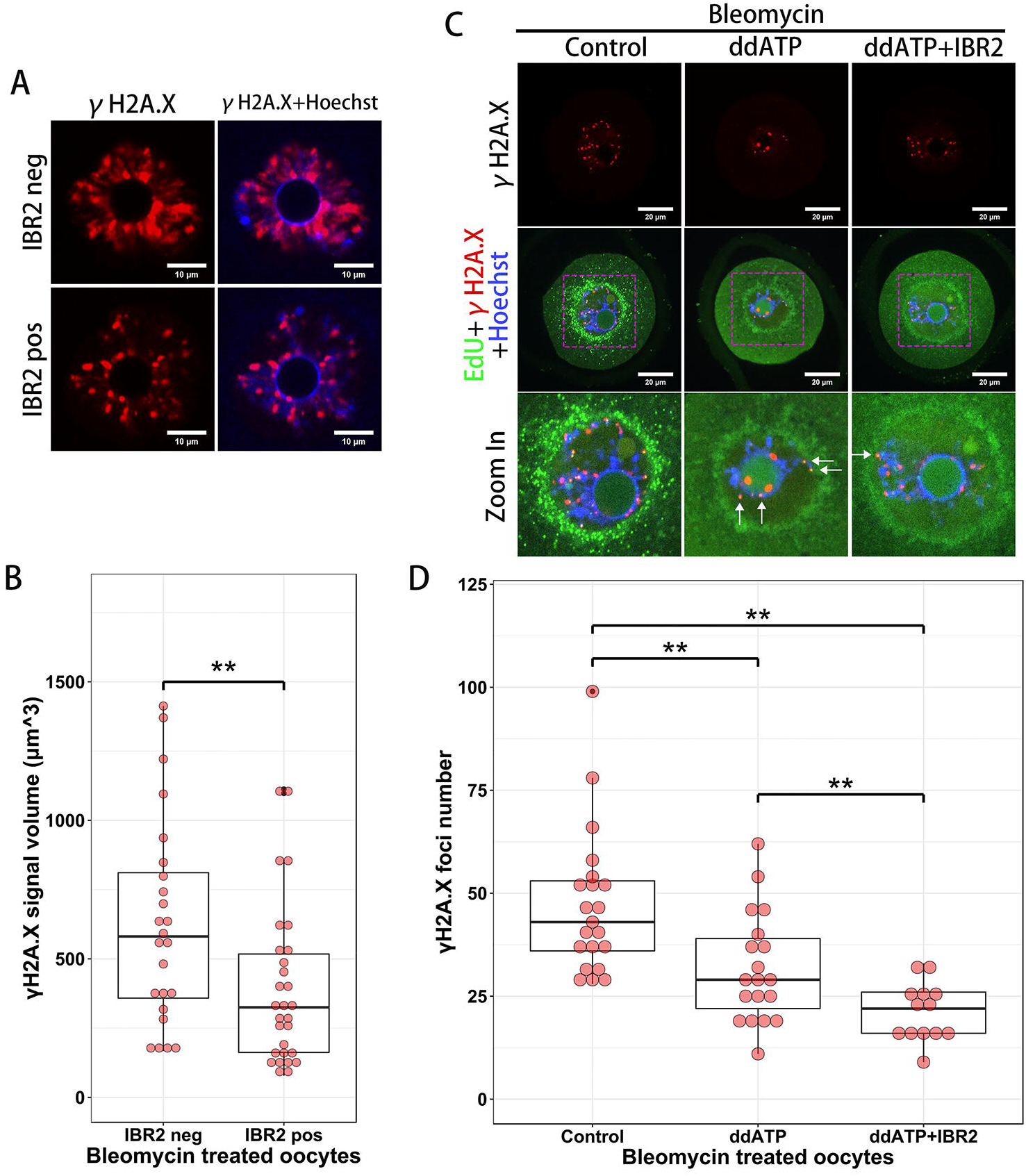
DNA damage has been amplified in SN oocytes. (**A** and **B**) After 5 hours treatment of 100 μM Rad51 inhibitor IBR2, oocytes are treated with 1 μM Bleomycin for 1 hour and released oocytes from Bleomycin for 4 hours. γH2A.X signals can been seen attenuated in IBR2 treated oocytes. (**C** and **D**) We treated oocytes with ddATP (100 μM) or with both ddATP and IBR2 (100 μM) for 5 hours and then treated oocytes with 0.5 μM Bleomycin for 1 hour and released oocytes from Bleomycin for 12 hours. Compared with control oocytes, γH2A.X foci numbers were less in ddATP group and further less in ddATP+IBR2 group. EdU signals in ddATP and ddATP+IBR2 group oocytes are marked by arrows.

As we had found Rad51 filaments exist beside to the newly synthesized DNA (Figure 2C), we want to know whether these Rad51 filaments left behind strand exchange could amplify the DSBs. To test it, we plan to inhibit the DNA replication in oocytes and examine whether the DSB number could be increased. At first, we used DNA polymerase inhibitor Aphidicolin to treat the oocytes, but we found Aphidicolin not only blocked the BIR in oocytes but also induce additional DSBs (Figure S2), indicating Aphidicolin is genotoxic to oocytes. So we next chose another DNA replication inhibitor ddATP which could block the mtDNA replication but could only delay the nuclear DNA replication. We treated oocytes with 100 μM ddATP for 5 hours and induced DSBs in oocyte with 0.5 μM Bleomycin for 1 hour, and then release the oocytes from Bleomycin for 12 hours. As a result we found ddATP could indeed suppress the replication of mtDNA but couldn’t fully suppress the nuclear DNA replication in oocytes (Figure 3C). But to our surprise, we found that ddATP treated DSB oocytes had fewer but not more γH2A.X foci than control oocytes (p < 0.01, Figure 3D), indicating nuclear DNA replication delay in SN oocytes suppressed the DNA damage amplification. In addition, when we treated DSB oocytes with both ddATP and IBR2, we found the γH2A.X foci number was further decreased comparing to the DSB oocytes only treated by ddATP (Figure 3D), indicating Rad51 and DNA replication played two independent roles in DNA damage amplification in SN oocytes.

### High DSB load induced BIR in NSN oocytes of hybrid mice

As BIR is mostly initiated by the invasion of the broken DNA end to a non-homologous sequence region, so we purposed that the DNA repair efficiency in oocytes might be affected by the different genome background. We then isolated oocytes from hybrid mice (filial generation of male C57 and female ICR mice, C57xICR F1) and oocytes from purebred ICR mice at the same age. Then we compared the γH2A.X foci numbers in ICR oocytes and C57xICR F1 oocytes 12 hours after releasing from Bleomycin treatment (1 μM for 1 hour). As a result we found the mean γH2A.X foci number in hybrid mouse oocytes was larger than that of purebred mouse oocytes (p < 0.01, Figure 4A and 4B), indicating that the sequence differences between homologous chromatids increased the DNA repair burden in SN oocytes.

**Figure 4.**
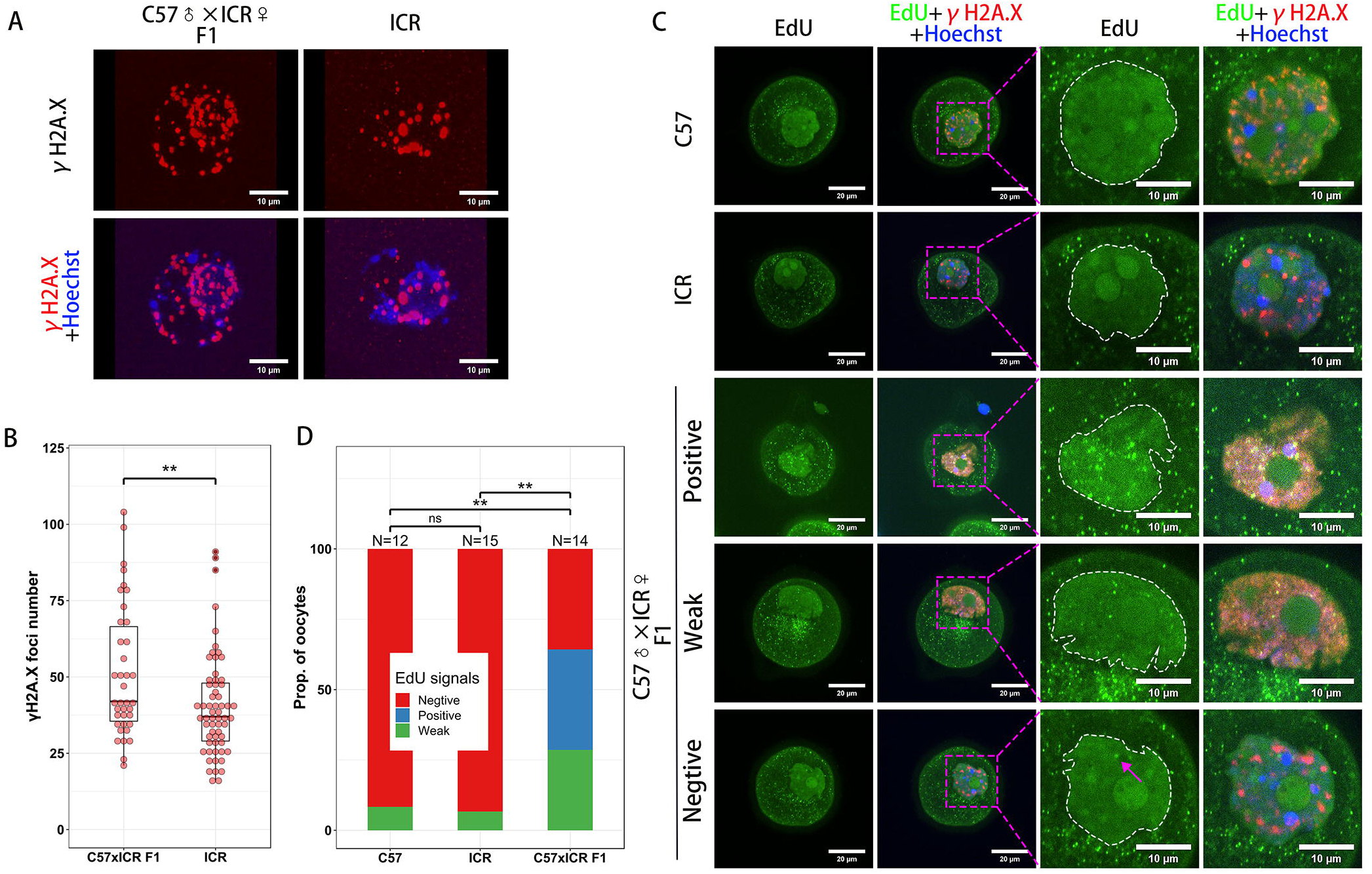
DNA damage repair is affected by the genome background in oocytes. (**A** and **B**) Hybrid mice (C57 x ICR F1) and purebred mice (ICR) oocytes were treated with 1 μM Bleomycin for 1 hour and released from Bleomycin for 12 hours. γH2A.X foci numbers in ICR oocytes were significantly less than C57 x ICR F1 oocytes. (**C** and **D**) Peri-antral follicular NSN oocytes of purebred mice (C57 and ICR) and hybrid mice (C57 x ICR F1) were treated with 40 μM Bleomycin for 1 hour and released from Bleomycin for 15 hours. EdU signals could be detected in hybrid NSN oocyte nuclear (enclosed by dashed line) but rare detected in purebred NSN oocyte nuclear.

To further analyze the effects of genome background on oocyte DNA repair, we treated the peri-antral follicular NSN oocytes from C57xICR F1, C57 and ICR mice with a higher dose of Bleomycin (40 μM for 1 hour) and release them for 15 hours. As a result we found BIR could be induced in the hybrid mouse oocytes more frequently than that in the purebred mouse oocytes (Figure 4C and 4D). This result further indicated sequence differences between homologous chromatids in oocytes would take effects on the DNA repair efficiency.

## Discussion

In this study, we used mouse oocytes as a model to analyze the DNA DSB repair in late G2 phase cells. We found DSBs in the SN oocytes, which have a condensed DNA configuration, but not in NSN oocytes could induce a type of small-scale BIRs. As the oocyte BIR induced EdU signal sizes were comparable with the mtDNA replication induced EdU signal sizes, and mouse mtDNA length ranges from 16299 to 16301 bp (Bayona-Bafaluy *et al.* 2003), so the scale of the length of BIR in SN oocytes should be approximately 10k to 20k bp. Evidences showed that the tract length of gene conversion is about 200–300 bp (Mansai *et al.* 2011) which is obviously less than the length of BIR mediated DNA synthesis in oocytes. On the other hand, these small-scale BIR in SN oocytes is also different with the classical BIR which would replicate DNA from breakpoint to the chromosome end or replicate a large genome fragment (Mancera *et al.* 2008; Ma *et al.* 2019b), so the small-scale BIRs in SN oocytes might be a new type of BIR.

In this study, we found the DSB induced EdU signals were generally adjacent to the Rad51 foci, indicating Rad51-binding broken DNA end hadn’t been fully exchanged with the template DNA. The partially strand exchange in SN oocytes might be caused by two reasons: the condensed DNA configuration or the sequence difference between the template and the broken DNA end. As micro-homology sequences had been detected in the breakpoints of most CGRs, so we speculated that the partially strand exchange in SN oocytes might be caused by the invasion of broken DNA ends to the micro-homology regions (Figure 5A).

**Figure 5.**
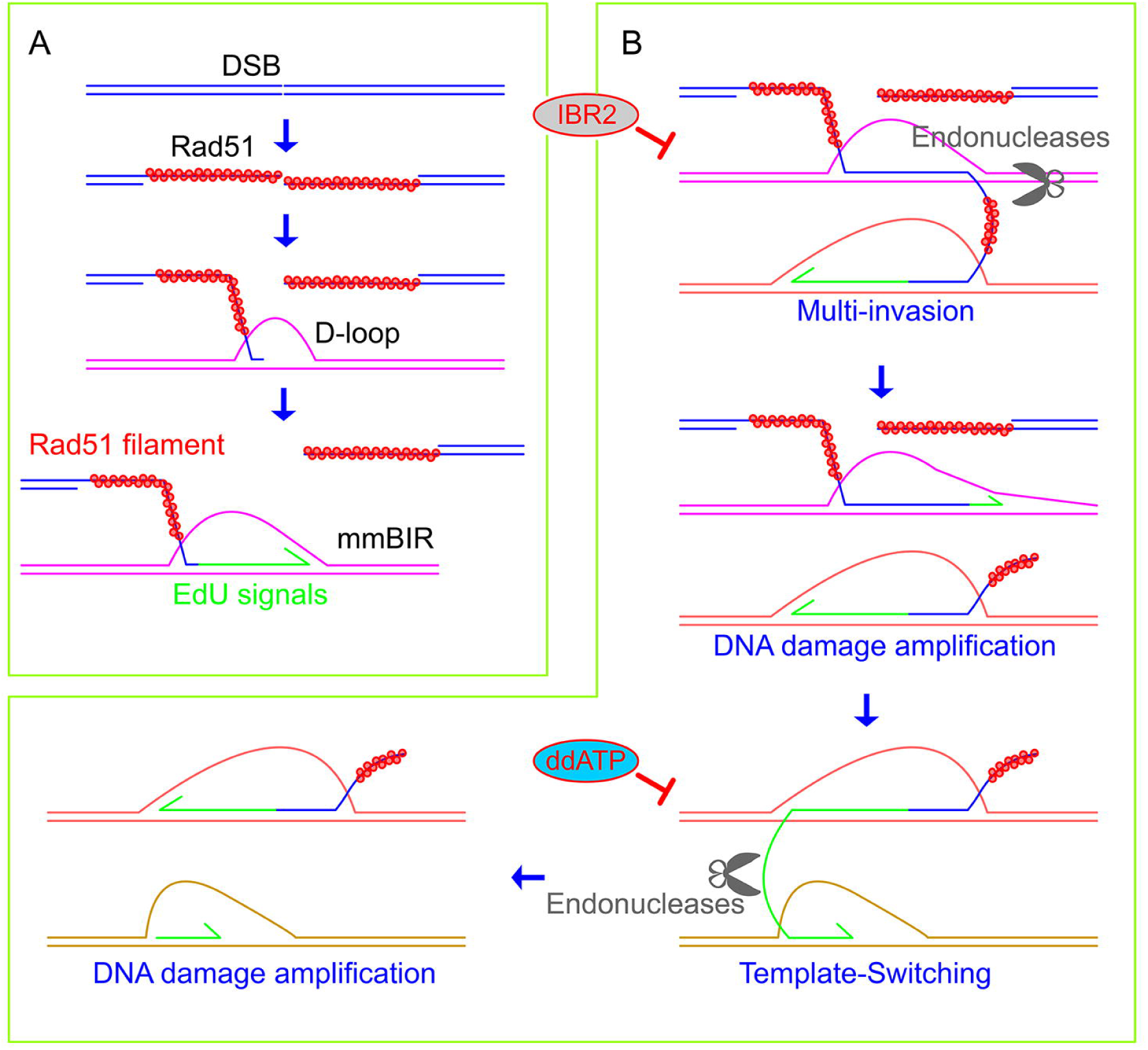
Supposed models of DNA DSB repair in SN oocytes. (**A**) Partially strand-exchange in SN oocytes make Rad51 filament exist beside the newly synthesized DNA. (**B**) The DNA damage in SN oocytes can be amplified by both Rad51 mediated multi-invasion and replication dependent template switching. Rad51 mediated multi-invasion can be suppressed by Rad51 inhibitor IBR2 whereas replication dependent template switching can be suppressed by ddATP.

Rad52 had been proven to mediated BIR in S phase of mammalian cells (Sotiriou *et al.* 2016), whereas Rad51 had been proven to be essential for the BIR in yeast (Davis and Symington 2004). In this study and our previous work (Ma *et al.* 2019a) we found Rad51 inhibitors RI-1 and IBR2 could suppress the BIR in SN oocytes, indicating BIR in late G2 phase cells was Rad51 dependent.

It had been reported that Aphidicolin and Hydroxyurea induced replication stress could induce the formation of rare CNVs in mammalian cells (Arlt *et al.* 2009; Arlt *et al.* 2011; Arlt *et al.* 2012). In this study we found that Aphidicolin could fully inhibit the BIR in SN oocytes, indicating DNA polymerase α and/or δ had participated in the DSB induced DNA replication. In oocytes, we found Aphidicolin could not only inhibit DNA replication but also induced additional DSBs, however, whether are these Aphidicolin induced DSBs associated with DNA polymerases is not known. It is possible that Aphidicolin induced DSBs in G2 phase are also causing factor of rare CNV formation.

Although we hadn’t found the direct evidence of Rad51 mediated broken DNA end multi-invasion, we indeed found the DSB numbers had been amplified in oocytes. We found the DSB amplification in oocytes could be suppressed by Rad51 inhibitor or by the DNA replication inhibitor ddATP. These results might suggest that the DSB amplification is not only induced by multi-invasion but also induced by DNA replication mediated template switching (Figure 5B). However, all these speculations need a further examination.

In our study we also compared the DNA repair difference between hybrid and purebred mouse oocytes. Compared to the purebred mouse oocytes, we found that the DSB repair in hybrid mouse SN oocytes was less efficiency. We also found higher DNA DSB load could induce BIR in hybrid mouse NSN oocytes. These results suggested that homologous chromatid could be used as template to repair the DSBs in oocytes and the sequence difference between homologous chromatids would affect the DNA repair efficiency and fidelity.

## Experimental procedures

### Oocytes isolation and *in vitro* culture

All of the animal experiments in this study were approved by the ethics committee of Guangdong Second Provincial General Hospital. Mice used in this study for oocyte collection were ICR, C57BL/6J (C57) and the first filial generation of female ICR and male C57 mice. If not mentioned, oocytes used in this study were isolated from ICR mice. The large antral follicle NSN oocytes and SN oocytes were isolated directly from ovaries of 8-12 weeks old female mice. The peri-antral follicles were isolated from ovaries of 8-12 weeks old female mice. To get peri-antral follicular oocytes, peri-antral follicles were treated with Trypsin-EDTA (25053CI, Corning) at 37□ for 25 minutes and oocytes were isolated from somatic cells with mouth pipette. To block the oocytes from meiosis resumption, all of the manipulations of oocytes in this study were under the M2 medium (Sigma, M7167) with 2.5 μM Milrinone (MCE, HY-14252).

### Treatment of oocytes by molecule compounds

DSBs in oocytes were introduced with Bleomycin at different concentrations (0.1, 0.5, 1, 10 or 40 μM) for 1 hour. To inhibit the activity of Rad51, we treated the oocytes with 100 μM Rad51 inhibitor IBR2 (MCE, HY-103710) whereas control oocytes were treated with DMSO. To inhibit the nuclear DNA polymerase activity, 2 μM Aphidicolin (MCE, HY-N6733) was used to treat the oocytes. To delay the nuclear DNA replication, 100 μM ddATP (Apexbio, B8136) was used to treat the oocytes.

### Immunofluorescence labeling

To immunofluorescence label the endogenous proteins in oocytes, oocytes were fixed with 4% Paraformaldehyde Fix Solution (Sangon, E672002) at room temperature (RT) for 15-30 minutes. Then in all following steps, the solutions used for immunofluorescence labeling were made up with PBST (0.1% Tween-20 in PBS). After fixation, oocytes were treated with 0.3% Triton X-100 at RT for 20 minutes. To unmask the antigen epitopes of specific endogenous proteins (Rad51 in this study), oocytes were treated with Quick Antigen Retrieval Solution for Frozen Sections (Beyotime, P0090) at RT for 40 minutes. Then oocytes were washed three times with PBST and blocked with 1% BSA at RT for 1 hour. Then oocytes were incubated with primary antibodies at 4 □ overnight and washed 5 times using PBST. After that oocytes were incubated with secondary antibodies at RT for 2-3 hours and washed 5 times with PBST. Then oocytes were stained with Hoechst for 1 hour and observed using the Andor live cell station system. The primary antibodies used in this experiment were: Rad51 antibodies (Abcam, ab133534; and Zen Bioscience, 200514), Mitofilin antibody (Proteintech, 10179-1-AP) and γH2A.X antibody (Bioworld, BS4760).

### EdU labeling

To label the new synthesized DNA in oocytes, the oocytes were cultured in M2 medium with 10 μM 5-ethynyl-2’-deoxyuridine (EdU, beyotime, ST067). Then oocytes were fixed with 4% Paraformaldehyde for 15 minutes and permeated with 0.3% Triton X-100 for 15 minutes. After incubation of primary and secondary antibodies, oocytes were treated with the click reaction buffer (beyotime, C0071S) at RT for 1 hour. Then oocytes were washed with PBST 5 times and stained with Hoechst.

### Statistic methods

The EdU signal size, γH2A.X foci number and γH2A.X foci volume of oocytes were measured or counted by the Fiji software (https://imagej.nih.gov). Students’ T test were used in this study for hypothesis test. P-value < 0.01 was recognized as very significant and marked with **; p-value < 0.05 and ≥ 0.01 was recognized as significant and marked with *. P-value ≿ 0.05 was recognized as not significant and marked with ‘ns’.

## Author contributions

J.Y.M. and X.H.O. conceived and planned the experiments. J.Y.M, X.F. and F.Y.X. carried out the experiments. J.Y.M., S.M.L., S.L. and L.N.C. contributed to sample preparation. J.Y.M. contributed to the interpretation of the results and took the lead in writing the manuscript. All authors provided critical feedback of the manuscript.

## Acknowledgements

We thank XY Fan, XH Sun and other members in Fertility Preservation Lab for their helps in this study.

---

**Figure S1. Cytoplasmic EdU signals colocalize with mitochondria in mouse oocytes.** Mitochondria is marked by mitofilin (red).

**Figure S2. Aphidicolin induces DSBs and inhibits BIR in oocytes.** (A) Aphidicolin (2 μM for 15 hours) induces DSBs in oocytes. (B) Aphidicolin (2 μM) blocked the BIR induced by Bleomycin (1 hour treatment of 1 μM Bleomycin for 1 hour and release oocytes from Bleomycin for 12 hours).

