## Supplementary figures and images for "Double-strand breaks can induce DNA replication and damage amplification in G2 phase-like oocytes of mice"

### Figure S1

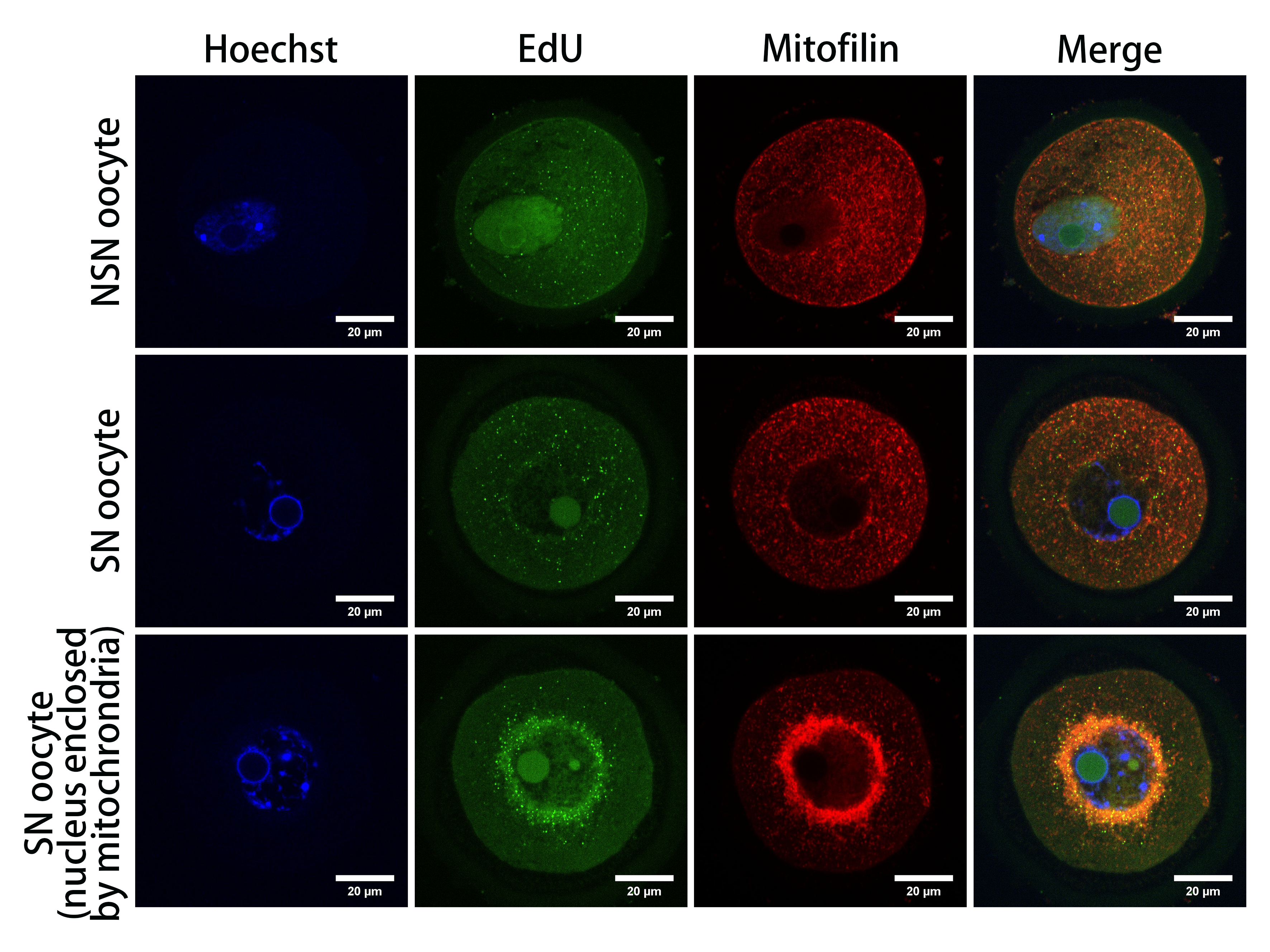

### Figure S2

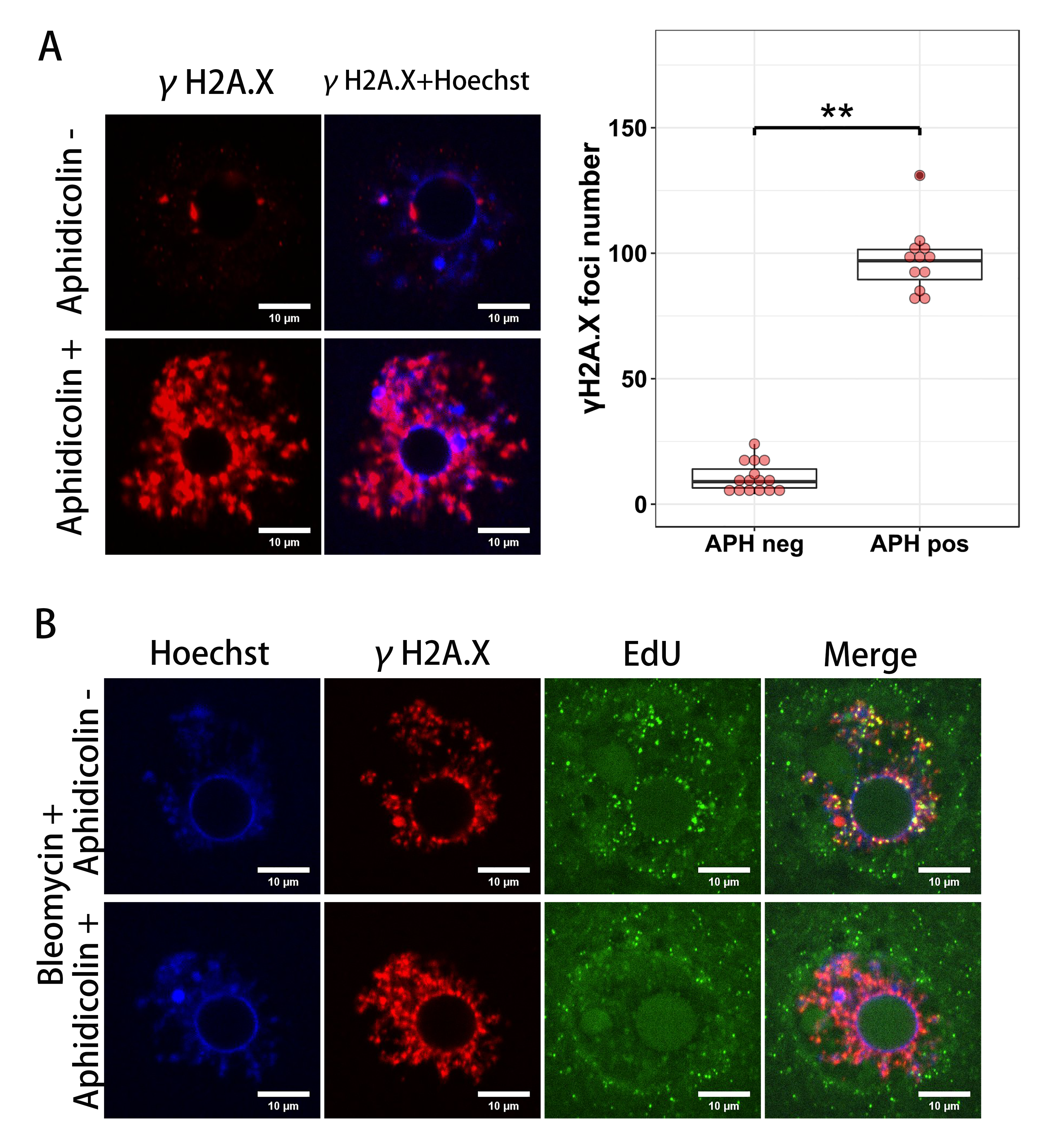
